# Genomewide Analyses of Psychological Resilience in US Army Soldiers

**DOI:** 10.1101/516716

**Authors:** Murray B. Stein, Karmel W. Choi, Sonia Jain, Laura Campbell-Sills, Chia-Yen Chen, Joel Gelernter, Feng He, Steven G. Heeringa, Adam X. Maihofer, Caroline M. Nievergelt, Matthew K. Nock, Stephan Ripke, Xiaoying Sun, Ronald C. Kessler, Jordan W. Smoller, Robert J. Ursano

## Abstract

Though a growing body of preclinical and translational research is illuminating a biological basis for resilience to stress, little is known about the genetic basis of psychological resilience in humans. We conducted genomewide association studies (GWAS) of self-assessed (by questionnaire) and outcome-based (incident mental disorders from pre- to post-deployment) resilience among European (EUR) ancestry soldiers in the Army Study To Assess Risk and Resilience in Servicemembers (STARRS). Self-assessed resilience (*N=11,492*) was found to have significant common-variant heritability (h^2^=0.162, se=0.050, p=5.37e-4), and to be significantly negatively genetically correlated with neuroticism (r_g_= −0.388, p=0.0092). GWAS results from the EUR soldiers revealed a genomewide significant locus (4 SNPs in LD; top SNP: rs4260523, p=5.654e-09) on an intergenic region on Chr 4 upstream from *DCLK2* (Doublecortin-Like Kinase 2), a member of the doublecortin (DCX) family of kinases that promote survival and regeneration of injured neurons. A second gene, *KLHL36* (Kelch Like Family Member 36) was detected at gene-wise genomewide significance (p=1.89e-06). A polygenic risk score derived from the self-assessed resilience GWAS was not significantly associated with outcome-based resilience. In very preliminary results, genomewide significant association with outcome-based resilience was found for one locus (top SNP: rs12580015) on Chr 12 downstream from *SLC15A5* (solute carrier family 15 member 5) in the small group (*N=*581) of subjects exposed to the highest level of deployment stress. The further study of genetic determinants of resilience has the potential to illuminate the molecular bases of stress-related psychopathology and potentially point to new avenues for therapeutic intervention.

## INTRODUCTION

Exposure to traumatic stressors is pervasive worldwide; in the United States, lifetime prevalence of a traumatic event is estimated at 70% [Benjet et al. 2016]. Individuals exposed to traumatic stressors are at heightened risk for psychiatric disorders including but not limited to posttraumatic stress disorder (PTSD) [Howlett and Stein 2016; Rosellini et al. 2018]. However, only a subset of individuals exposed to traumatic stressors subsequently develops such disorders, indicating that many can be considered resilient to those effects on psychopathology [Galatzer-Levy et al. 2018; Kalisch et al. 2014] While varying definitions exist in the literature, most conceptualize psychological resilience as successful adaptation in the face of adversity — often facilitated by personality traits or other individual differences [Kalisch et al. 2017; Pietrzak et al. 2014], and reflected in the absence of negative mental health outcomes where otherwise expected [Bonanno et al. 2011; Southwick and Charney 2012].

Though a growing body of preclinical and translational research is illuminating biological mechanisms of stress resilience [McEwen et al. 2015], relatively little is known about the genetic basis of psychological resilience in humans [Feder et al. 2018]. Twin studies have suggested that self-(or parent-) assessed resilience – defined as a perceived capacity to cope adaptively with stressors – is moderately heritable (∼30-50%) [Amstadter et al. 2014; Waaktaar and Torgersen 2012; Wolf et al. 2018]. Studies in twin samples and unrelated individuals have also suggested that other traits reflecting positive psychological adjustment, such as subjective well-being and positive affect are partially heritable [Haworth et al. 2016; Rietveld et al. 2013; Wingo et al. 2017]. Notably, these heritable traits have also been associated with resilient outcomes following various stressors; for example, positive affect has been found to be protective against psychiatric symptoms following major disasters [Fredrickson et al. 2003] daily stressors [Ong et al. 2006] and chronic illness [Zautra et al. 2005].

To date, there have been a limited number of genetic studies of psychological resilience, with most of these investigating candidate genes (e.g., *SLC6A4\**5HTTLPR) [Stein et al. 2009] for what is certainly a highly polygenic trait and, often focusing exclusively on PTSD as the outcome (e.g., APOE epsilon4, or, nitric oxide pathway genes) [Bruenig et al. 2017; Mota et al. 2018]. One recent study examined self-reported resilience along with polygenic risk for depression in relation to major depression, finding additive effects, consistent with the notion that psychological characteristics associated with self-assessed resilience can be considered a buffer against stress.[Navrady et al. 2017]. Several other studies have examined polygenic risk scores for major depression as predictors of depression following life stressors [Colodro-Conde et al. 2017; Domingue et al. 2017]. But, to the best of our knowledge, no prior study has sought to identify genomewide variation associated with resilience as either a self-reported trait, or as an outcome following stress.

Using data from the Army Study To Assess Risk and Resilience in Servicemembers (STARRS), the aim of the present study is to use genome-wide association methods to identify genetic variants associated with resilience phenotypes, both as a self-assessed trait and as an empirically and prospectively defined outcome. For the former phenotype, we use a 5-item measure of self-assessed resilience, which we have shown in STARRS has protective associations with prospective mental health outcomes in deployed soldiers [Campbell-Sills et al. 2018]. Specifically, we found that greater pre-deployment self-assessed resilience was associated with decreased incidence of emotional disorder (AOR = 0.91; 95% CI = 0.84–0.98; P = .016) and increased odds of improved coping (AOR = 1.36; 95% CI = 1.24–1.49; P < .0005) after deployment. For the prospectively defined resilience phenotype, we use a prospectively determined composite mental health outcome following an index deployment to Afghanistan. We also determine the common-variant heritability of resilience in this generally young and mostly male sample, and explore its genetic correlations with several other mental and physical health-related phenotypes [Zheng et al. 2017]. We focus our analyses on soldiers of European ancestry, the largest group in STARRS, and the only ancestral group with out-of-sample publicly available GWAS data for estimating genetic correlations. Findings are expected to provide insight into the biological bases of psychological resilience.

## METHODS

### Subjects

Information in detail about the design and methodology of STARRS can be obtained in our prior report [Ursano et al. 2014]. Each of the participating institutions approved the human subjects and data protection procedures used in the study. As described below, the analyses presented here involved two large study components of STARRS.

#### New Soldier Study (NSS)

New soldiers took part in the NSS at the beginning of their basic training, which took place between April 2011 and November 2012 at one of three Army installations. Soldiers completed a computerized self-administered questionnaire (SAQ, described below) and 83.2% gave blood samples for DNA. Genotyping was conducted in samples from the first half of the cohort (NSS1; *N* = 7,999) and on a smaller subset of the second half of the cohort (NSS2; *N* = 2,835) (see Supplementary Materials for details). Data from subjects of European (EUR) ancestry in NSS1 (*N* = 4,750) and NSS2 (*N* = 1,817) were included in these GWAS meta-analysis of self-assessed resilience and in the subsequent derivation of a polygenic risk score (PRS) for self-assessed resilience (**Figure 1**).

**Figure 1.**
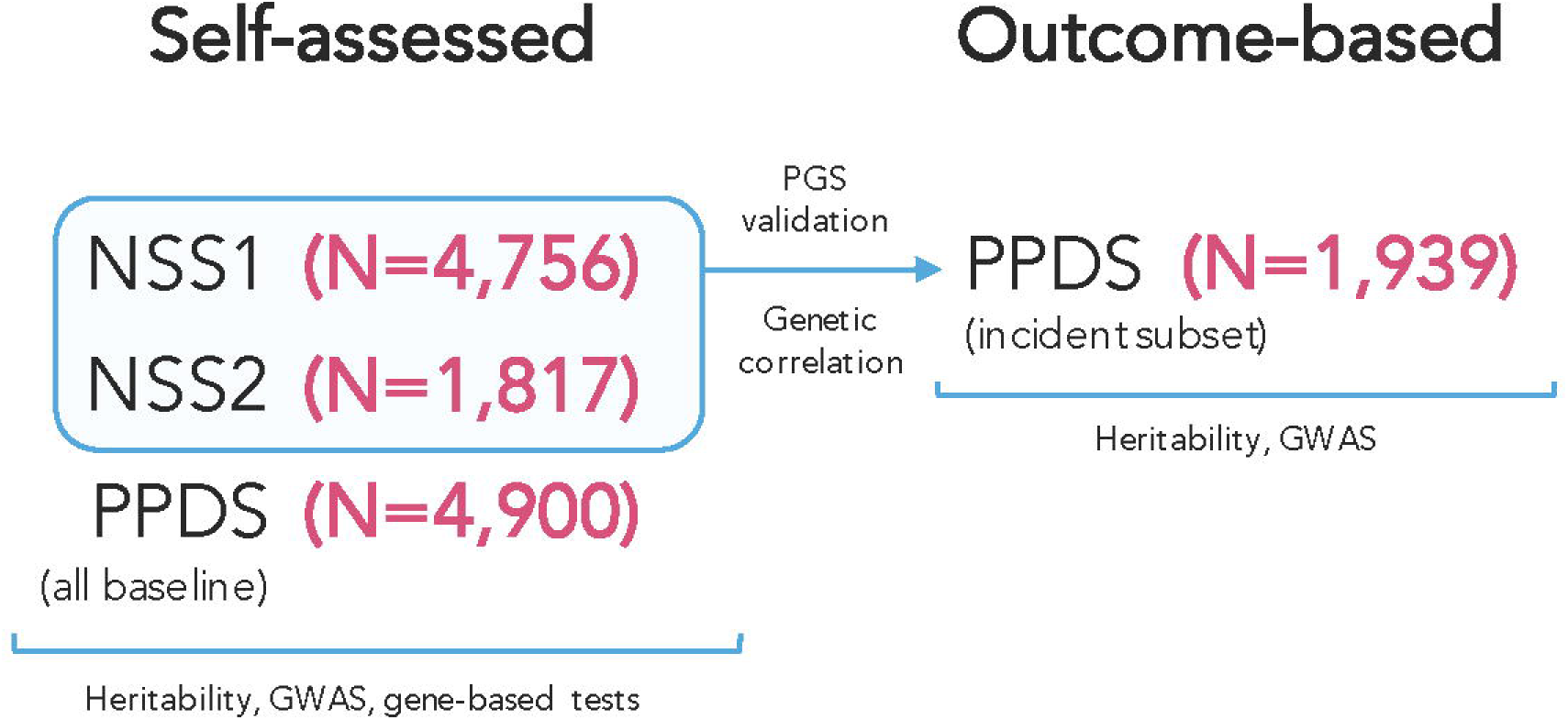
Cohorts Used for Analysis of Self-assessed and Outcome-based Resilience.

#### Pre/Post Deployment Study (PPDS)

US Army soldiers from three Brigade Combat Teams (BCTs) participated in the PPDS (*N* = 7,927 eligible soldiers were genotyped) that began in the first quarter of 2012. The data included in this report were collected at baseline (T0) 4-6 weeks prior to deployment to Afghanistan, and approximately 3- and 9-months following return from deployment. Data from EUR PPDS soldiers were included in the GWAS meta-analysis of self-assessed resilience and also in a GWAS of outcome-based resilience. Data from PPDS soldiers were <u>not</u> included, however, in the polygenic score (PRS) of self-assessed resilience that was derived in NSS1+NSS2 and subsequently tested in PPDS (i.e., they were independent) (**Figure 1**).

### Measures

#### Self-assessed resilience

Self-assessed resilience was measured using a STARRS 5-item self-report questionnaire that asked respondents to rate their ability to handle stress in various ways. Examples included “Keep calm and think of the right thing to do in a crisis” and “Keep your sense of humor in tense situations,” each rated 0 (poor) to 4 (excellent), and summed to yield a total resilience score ranging from 0-20. This STARRS self-report questionnaire has been found to have a unidimensional structure, demonstrates good internal consistency and, as noted above, has been shown to have predictive validity for resilient outcomes following exposure to deployment stress [Campbell-Sills et al. 2018].

#### Deployment (Combat) stress

Combat/deployment stress was quantified using a Deployment Stress Scale (DSS; theoretical range=0-16) used in our prior research with these cohorts [Campbell-Sills et al. 2018; Stein et al. 2015]. Higher DSS scores reflect greater exposure to traumatic deployment experiences, such as firing at the enemy/taking enemy fire or being exposed to severely wounded or dying people.

#### Outcome-based resilience

The Composite International Diagnostic Interview screening scales (CIDI-SC) [Kessler and Ustun 2004] were used to assess criteria for four common stress-related psychiatric disorders: major depression, generalized anxiety disorder, posttraumatic stress disorder and panic disorder. To assess new-onset, or incident disorders following deployment, our analytic sample was constrained to PPDS soldiers who met current criteria for *none* of these disorders pre-deployment (*N* = 1,939). Outcome-based resilience was defined as not meeting criteria for *any* of these incident disorders post-deployment.

### DNA Genotyping and Imputation

Detailed information on genotyping, genotype imputation, population assignment and principal component analysis for population stratification adjustment are included in our previous report [Stein et al. 2016] and in Supplementary Materials. Briefly, whole blood samples were shipped to Rutgers University Cell & DNA Repository (RUCDR), where they were frozen for later DNA extraction using standard methods. NSS1 and PPDS samples were genotyped using the Illumina OmniExpress + Exome array with additional custom content (*N* SNP = 967,537). NSS2 samples were genotyped on the Illumina PsychChip (*N* SNP = 571,054; 477,757 SNPs overlap with OmniExpress + Exome array).

Relatedness testing was carried out with PLINK v1.90 [Chang et al. 2015; Purcell et al. 2007] and pairs of subjects with π of >0.2 were identified, randomly retaining one member of each relative pair. We used a two-step pre-phasing/imputation approach for genotype imputation, with reference to the 1000 Genomes Project multi-ethnic panel (August 2012 phase 1 integrated release; 2,186 phased haplotypes with 40,318,245 variants). We removed SNPs that were not present in the 1000 Genomes Project reference panel, had non-matching alleles to 1000 Genome Project reference, or had ambiguous, unresolvable alleles (AT/GC SNPs with minor allele frequency [MAF] > 0.1). For the Illumina OmniExpress array 664,457 SNPs and for the Illumina PsychChip 360,704 SNPs entered the imputation procedure.

### Ancestry Assignment and Population Stratification Adjustment

Given the ancestral heterogeneity of the STARRS subjects, samples were assigned into major population groups (European [EUR], African, Latino or Asian). In order to avoid long range LD structure from interfering with the PCA analysis, we excluded SNPs in the MHC region (Chr 6:25-35Mb) and Chr 8 inversion (Chr 8:7-13Mb). Principal components (PCs) within each population group were then obtained for further population stratification adjustment. Details of these procedures are described in an earlier STARRS publication [Stein et al. 2016]. As noted above, results reported here are limited to the largest population group in the study, those of EUR descent.

### Genomic and Sample Quality Control (QC)

For QC purposes we kept autosomal SNPs with missing rate < 0.05; kept samples with individual-wise missing rate < 0.02; and kept SNPs with missing rate < 0.02. After QC, we merged our study samples with HapMap3 samples. We kept SNPs with minor allele frequency (MAF) > 0.05 and LD pruned at R^2^ > 0.05.

### Statistical Analysis

As noted above, analyses were limited to soldiers of EUR ancestry. First, we estimated the proportion of variance in self-assessed resilience and outcome-based resilience explained by common SNPs (i.e., SNP-heritability, *h*^*2*^_*g*_) with linear mixed models implemented in GCTA software [Yang et al. 2011].

Second, we used PLINK v1.90 [Chang et al. 2015; Purcell et al. 2007] with imputed SNP dosages to conduct genome-wide association tests for each type of resilience using linear regression (for self-reported resilience) and logistic regression (for dichotomized outcome-based resilience), each adjusted for age, sex, and the top 10 within-population PCs. We filtered out SNPs with MAF < 0.01 or imputation quality score (INFO) < 0.6, and performed HWE tests for the top SNPs from the association analysis. GWAS for self-assessed resilience was conducted in the three studies (NSS1, NSS2 and PPDS) separately and then meta-analyzed across studies (**Figure 1**). Meta-analysis was conducted using an inverse variance-weighted fixed effects model in PLINK. Again, GWAS for outcome-based resilience was conducted in the PPDS, exclusively among soldiers with no disorder prior to the index deployment (**Figure 1**). A p-value < 5 × 10^−8^ was used as the threshold for genome-wide significance whereas results at p-value < 1 × 10^−6^ are reported as genome-wide suggestive.

To follow up on GWAS results for self-assessed resilience, we performed gene-based tests using software MAGMA [de Leeuw et al. 2015] within the FUMA suite [Watanabe et al. 2017]. (These analyses were not conducted for outcome-based resilience, given the small sample size available for that phenotype.) The gene-based test in MAGMA provides association tests for each gene (i.e., genome-wide gene-association study [GWGAS]; N = 18,167 protein coding genes) by aggregating SNPs within the gene region. We used the final meta-analytic results and the 1000 Genomes Project European LD reference for this analysis. For the gene-based analysis, we used a combined mean and top SNP association model; the significance level after Bonferroni correction is 0.05/18167= 2.75×10^−6^.

Polygenic risk scores (PRS) [Euesden et al. 2015] for self-assessed resilience were constructed using summary statistics from the NSS1/NSS2 GWAS data only, and applied to PPDS. After removal of ambiguous SNPs, we clumped summary statistics to limit inclusion of highly correlated SNPs, using a linkage disequilibrium r^2^ of 0.25 to select index SNPs within each 250kb window. Clumped summary statistics were used to compute PRS from our genomic data that included SNPs whose effect sizes met the following p-value thresholds, in decreasing order of stringency: <0.001, 0.01, 0.05, 0.10, 0.50, and 1.0. PRS were calculated as the total sum of risk alleles at each eligible SNP weighted by their estimated effect size, divided by total number of SNPs included for scoring.

We used LD Score Regression (LDSC) [Bulik-Sullivan et al. 2015] implemented on LD Hub (http://ldsc.broadinstitute.org) [Zheng et al. 2017] referencing publicly available meta-analytic GWAS results to test genetic correlations between self-assessed resilience and five traits of theoretical relevance to resilience: major depression (a disorder frequently studied as an outcome in prior resilience studies) [Major Depressive Disorder Working Group of the Psychiatric et al. 2013], neuroticism (a personality trait frequently associated with poor resilience), subjective well-being [Okbay et al. 2016], intelligence [Sniekers et al. 2017] and hippocampal volume [Hibar et al. 2015]

## RESULTS

### Sample Descriptions

For self-assessed resilience, the sex, age, marital status, and education composition of our analyzed participants along with average resilience scores are shown in **Table 1**. For outcome-based resilience, 80.4% (*N* = 1558) of the PPDS soldiers eligible for analysis were resilient post-deployment, whereas 19.7% (*N* = 381) had developed an incident deployment-related mental disorder.

**Table 1.** Study participants with self-assessed resilience scores, and sex and age distributions in the samples

| Self-Assessed Resilience |  |  |  |  |
| --- | --- | --- | --- | --- |
| <u>Study</u> | <u>Ancestry</u> | <u>N</u> | <u>Mean</u> | <u>SD</u> |
| NSS 1 | EUR | 4756 | 13.57 | 4.31 |
| NSS 2 | EUR | 1817 | 13.41 | 4.47 |
| PPDS | EUR | 4900 | 14.75 | 4.23 |

| Sociodemographic Characteristics |  |  |  |
| --- | --- | --- | --- |
|  | NSS1 | NSS2 | PPDS |
| Sex (% Male) | 81.4% | 77.8% | 92.8% |
| Age Year (Mean [SD]) | 21.0 (3.3) | 20.3 (3.2) | 25.9 (5.9) |
| Marital Status (% Ever Married) | 12.0% | 9.1% % | 54.0% |
| Education (% >= High School) | 88.7% | 90.7% | 92.8% |

### Genome-wide Association Analyses of Self-Assessed Resilience

In the meta-analysis of EUR ancestry GWASs across the three cohorts (NSS1, NSS2, and PPDS), we identified 4 genome-wide significant SNPs on Chr4 (reflecting one genomewide significant locus; lead SNP rs4260523, beta = 0.352, p = 5.65×10^−9^) in an intergenic region upstream from *DCLK2* (see **Figure 2** for Manhattan plot [lambda = 1.03] and **Figure 3** for regional plot). These and two other independent genome-wide suggestive (p < 10^−6^) loci are shown in **Supplementary Table S1**.

**Figure 2.**
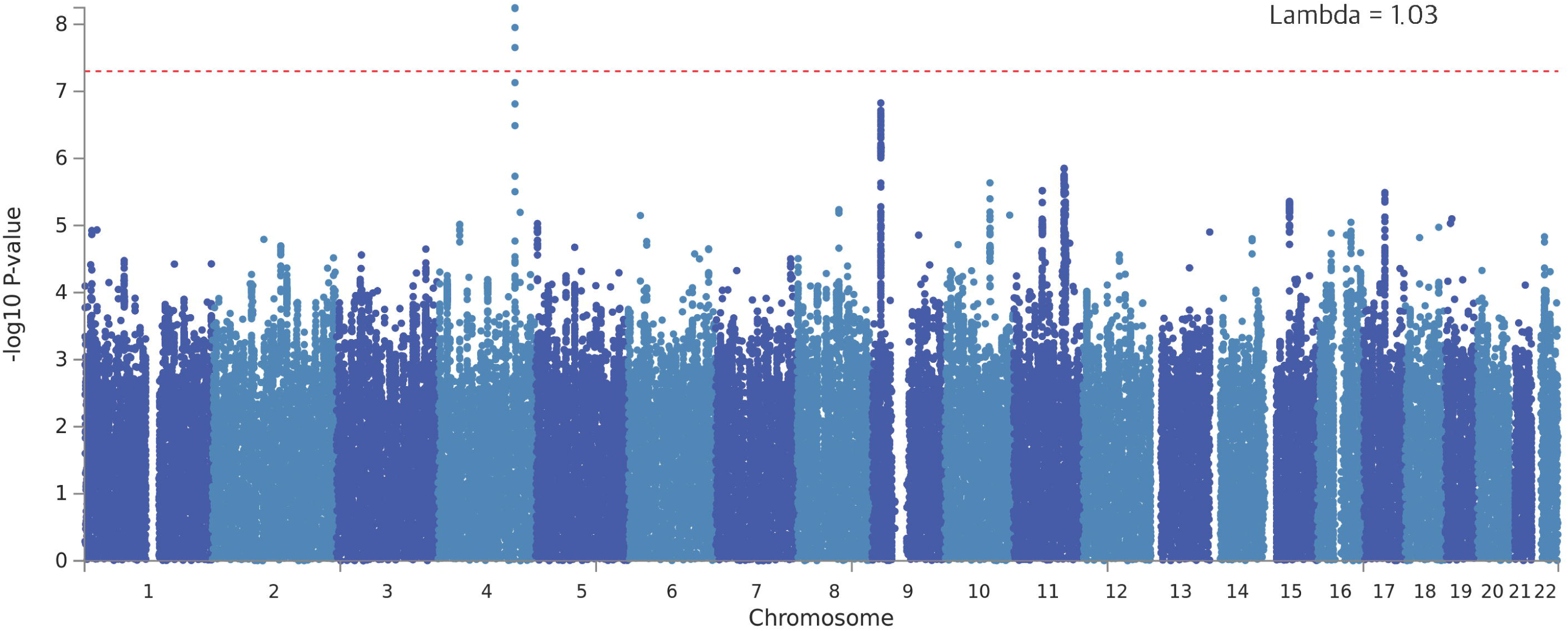
Manhattan plot of NSS1, NSS2, and PPDS Self-Assessed Resilience Genome-Wide Association Study (GWAS) in Soldiers of European (EUR) Ancestry.

**Figure 3.**
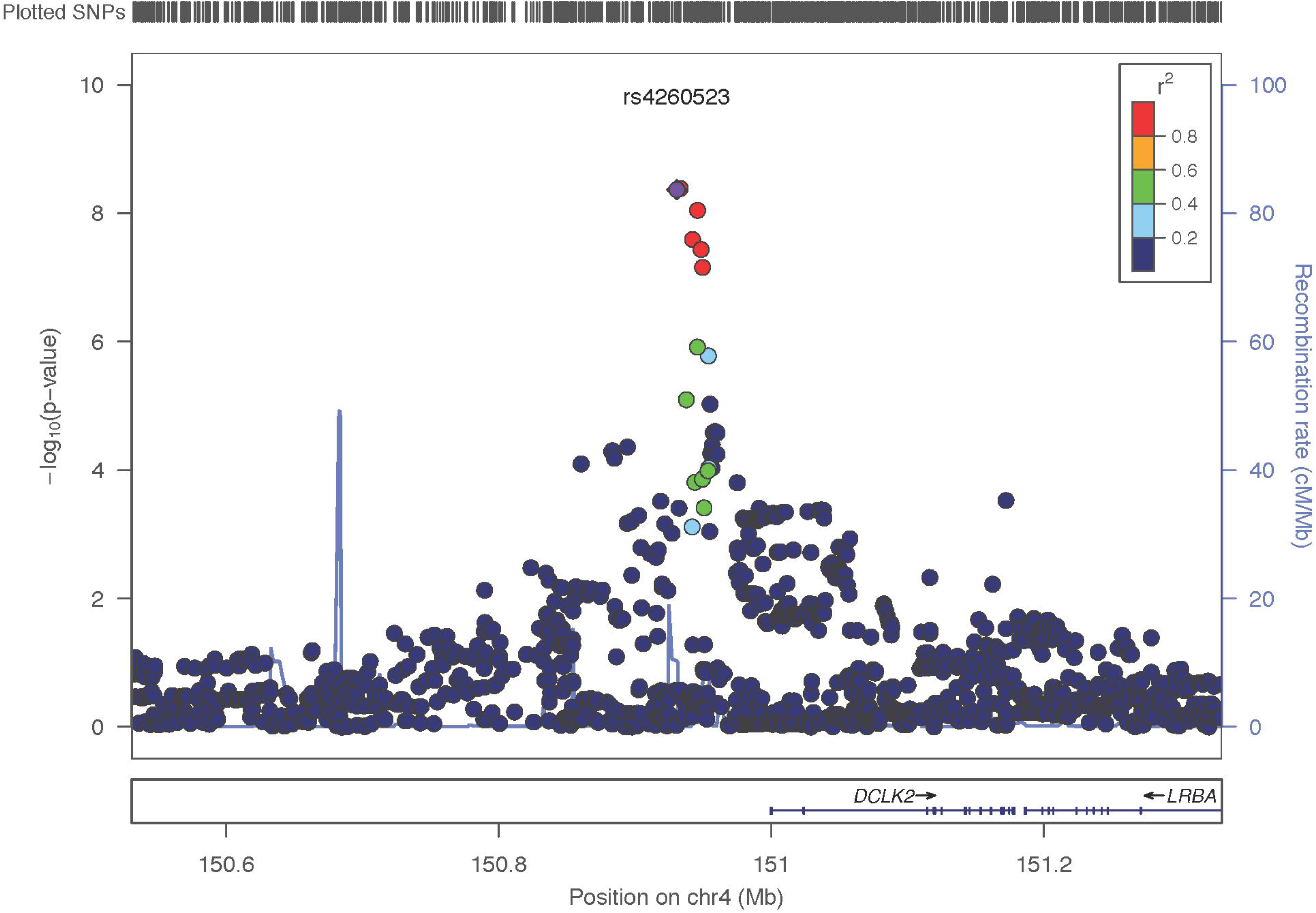
Locus-zoom plot showing region on Chr 4 containing the genome-wide significant markers in the NSS1, NSS2, and PPDS Self-Assessed Resilience EUR GWAS.

#### Genome-wide Gene Association Analysis (GWGAS) of Self-Assessed Resilience

There was one significant gene in the self-assessed resilience meta-analysis, identified via genome-wide gene-association study (GWGAS; **Supplemental Figure S1**) with MAGMA after Bonferroni correction: *KLHL36* (Kelch Like Family Member 36; gene ID 79786), on chromosome 16, with a p-value = 1.89×10^−6^ obtained by aggregating 134 SNPs in the region. We list all the genes in the GWGAS and highlight the top 6 genes with the most significant p-values (< 10^−4^) from the EUR meta-analysis in **Supplementary Table S2**.

#### SNP-based heritability of Self-Assessed Resilience

Using GCTA [Yang et al. 2011] we estimated SNP-based heritability of self-assessed resilience in the EUR subjects (N = 9,932) to be h^2^_g_ = 0.162, se = 0.050, p = 5.37×10^−4^.

#### Genetic Correlations of Self-Assessed Resilience with Other Traits

Using LDSC as implemented in LDHub we found, after Bonferroni correction (p < 0.01, accounting for 5 comparisons) one statistically significant genetic correlation between self-assessed resilience and the 5 selected relevant traits of *a priori* interest. We observed a significant (negative) genetic correlation with neuroticism (from UK Biobank) (r_g_ = −0.388, p = 0.0092), but not with the other 4 traits including major depressive disorder (r_g_ = −0.464, p = 0.077), subjective well-being (r_g_ = 0.269, p = 0.083), intelligence (r_g_ = −0.071, p = 0.579) or hippocampal volume (r_g_ = −0.223, p = 0.463).

#### Polygenic Risk Scores for Self-Assessed Resilience Related to Outcome-Based Resilience

PRS derived from self-assessed resilience in EUR NSS1+NSS2 were not significantly associated with outcome-based resilience in EUR PPDS at any tested p-value level (**Supplementary Figure S3**), though all were associated with nominally higher odds for outcome-based resilience.

### Genome-wide Association Analyses of Outcome-Based Resilience

In our exploratory (given the small sample size) GWAS of outcome-based resilience that included all eligible deployed soldiers (*N* = 1,939) we did not observe any genomewide significant SNPs (**Supplementary Table S3a**), even when adjusting for individual levels of deployment stress exposure (**Supplementary Table S3b)**. When we restricted analysis only to soldiers (*N* = 581) who had experienced high deployment stress exposure (deployment stress score >=8 out of a possible 16) we found one genomewide significant locus associated with outcome-based resilience (top SNP: rs12580015*C, OR = 0.42, p = 2.37×10^8^) in LOC101928362, less than 0.1 Mb downstream from *SLC15A5* (solute carrier family 15 member 5; gene ID: 729025) on Chr 12p12.3; (**Supplemental Figure 2a** [Manhattan Plot] and **Supplemental Figure 2b** [Regional Plot] and **Supplemental Table S3c**). SNP-based heritability of outcomes based resilience in the EUR subjects (N = 1,853) was not statistically significant. There was no overlap in the genomewide significant or suggestive (p < 10^−6^) SNPs associated with self-assessed and outcome-based resilience (in either the full eligible sample or the high combat stress exposure group).

Finally, we calculated the genetic correlation (r_g_) between self-assessed resilience in NSS1+NSS2 and outcome-based resilience in PPDS. Although the magnitude of the correlation and its positive directionality were consistent with expectations, the r_g_ estimate of 0.663 (se 0.422) was not statistically significant (p = 0.123), likely reflecting the very small sample size available for the outcome-based phenotype.

## DISCUSSION

Identifying factors that contribute to psychological resilience in the face of stressors is of paramount importance to the understanding of mental health and wellbeing. Several recent reviews have pointed to a multitude of neurobiological factors believed to play a role in resilience [Feder et al. 2018; Menard et al. 2016; Pfau and Russo 2015] including diverse stress response systems [McEwen et al. 2015]. While the potential genetic underpinnings of these factors have begun to receive attention, studies to date have focused on candidate gene (or epigenetic) [Binder 2017] involvement [Feder et al. 2018; McEwen 2016; Menard et al. 2016]. Here, we report results from what we believe to be the first GWAS of psychological resilience, and have done so in military population-based samples. Consistent with twin studies we find strong evidence that self-assessed resilience has a heritable basis (SNP-based heritability 16%) in this population. We also find a strong negative genetic correlation between self-assessed resilience and a personality trait known to be a risk factor for psychopathology, neuroticism. And we discover preliminary associations between several specific genes and self-assessed (*DCLK2 and KLHL36*) resilience.

*DCLK2* is an intracellular enzyme preferentially expressed in the brain and particularly enriched in cerebral cortex and hippocampus (www.proteinatlas.org/) [Uhlen et al. 2015]. Mice lacking *DCLK2* have altered hippocampal development and spontaneous seizures [Kerjan et al. 2009]. Members of the doublecortin (DCX) family of kinases promote survival and regeneration of injured neurons [Nawabi et al. 2015]. Genetic variations in *DCX* genes including deletions, nonsense, frameshift and missense mutations have been associated with lissencephaly (characterized by the absence of normal convolutions in the cerebral cortex and microcephaly). Certain types of genetic variation in *DCLK2* might therefore be associated with less deleterious changes in brain structure or cognitive function that could influence resilience. *DCLK2* is a neighboring gene to *NR3C2* (a mineralicorticoid receptor gene associated in one study with stress resilience [ter Heegde et al. 2015]) and we considered the possibility that SNPs we identified as being in an intergenic region of *DCLK2* might regulate expression of *NR3C2*. But according to GTeX v7 none of the SNPs in that region (see Table S1) of Chr4 were labeled as eQTLs in any tissue. A SNP in *DCLK2* (rs11947645, approximately 0.4 MB downstream from our top SNP) was observed to be the top hit (though below genomewide significance at p = 1.47×10^−06^) in a GWAS of social skills (considered in that study to be an autistic-like trait) in a population-based study of young adults [Jones et al. 2013]. Given the importance of strong social connectedness as a factor in resilience, one could imagine how being at genetic risk for poor social skills could result in lower resilience to stressors.

*KLHL36* emerged in association with self-assessed resilience in the gene-based analysis. The product of this gene ubiquinates proteins as part of their degradation pathway and is widely expressed in virtually all tissues. A SNP in *KLHL36* (rs12716755) has been reported to be a risk variant for late onset Alzheimer’s Disease. These observations and their implications for illuminating a role for *DCLK2* and *KLHL36* in resilience remain to be determined.

The importance of looking at prospectively defined outcomes in resilience research has recently been highlighted [Chmitorz et al. 2018]. While sample size was limited, we had the unique opportunity to explore genetic contributions to resilience in a prospective cohort where exposure to trauma was empirically measured. Our finding that a genomewide significant locus for outcomes-based resilience became visible only when restricting the analysis to those soldiers who had experienced the most combat stress exposure highlights the value of studying resilience in the context of stressful experiences. However, although ours is, to the best of our knowledge, the first study to include a prospectively determined cohort to assess resilience in a genomewide analysis, our sample size for that analysis was so small (N = 581 for the high-deployment stress exposed subgroup) that our observations must be considered more of a proof-of-feasibility than a discovery of risk-related variants. As such, we consider the association with *SLC15A5* to be preliminary, quite possibly a false-positive, and definitely in need of replication. We also found that polygenic scores for self-assessed resilience from NSS did not predict outcomes-based resilience in PPDS and that genetic correlation between the two traits was not statistically significant. These observations may signal that these two indicators of resilience – though linked at the phenotypic level [Campbell-Sills et al. 2018] – are relatively genetically distinct and may be related through environmental factors, although we cannot exclude the strong possibility that this null finding is because our samples were underpowered to detect a genetic correlation.

Our results should also be interpreted in light of several additional limitations. First and foremost, our study looks at prospectively determined resilience through the rather narrow lens of not developing a mental disorder during a stressful life period. As mentioned above, many other definitions of resilience could have been considered, but we were limited by the data at hand in our survey. Second, power to detect loci of modest effect is limited given our current sample sizes, and the precision of our effect sizes may be reduced given that resilience was studied here as a secondary trait [Yung and Lin 2016]. Third, since over 80% of our sample is comprised of men, all of European descent, our results may not generalize well to women or to other ancestry groups. Fourth, although we used a measure of self-reported resilience that, in our prior work, was shown to predict outcomes-based resilience in these cohorts [Campbell-Sills et al. 2018], it is not a well-studied, widely used measure of self-reported resilience such as the Connor-Davidson Resilience Scale [Connor and Davidson 2003] and variants thereof [Campbell-Sills and Stein 2007], and its relationship to other correlates of resilience such as positive affect is not currently known. Fifth, focused as we were on genetic risk factors, we did not test more complicated models that might have adjusted for other known experiential resilience risk factors such as childhood maltreatment, or other types of trauma. Such analyses will require much larger sample sizes able to accommodate multiple covariates and their interactions.

In summary, this set of genome-wide association studies confirms a genetic basis for self-assessed resilience, offers some insights into the possible molecular biological bases for resilience to stressors, and provides proof-of-concept that genomewide studies of outcomes-based resilience will be possible given adequate sample size. Greater exploration of the genetic bases of resilience – focused on variants that contribute to health, rather than disease [Schwartz et al. 2017] – will not only contribute to our understanding of the structure of psychopathology [Smoller et al. 2018] but may also identify actionable targets in the quest for precision psychiatry [Stein and Smoller 2018].

## Acknowledgments

The Army STARRS Team consists of:

Co-Principal Investigators: Robert J. Ursano, MD (Uniformed Services University of the Health Sciences) and Murray B. Stein, MD, MPH (University of California San Diego and VA San Diego Healthcare System)

Site Principal Investigators: Steven Heeringa, PhD (University of Michigan), James Wagner, PhD (University of Michigan) and Ronald C. Kessler, PhD (Harvard Medical School) Army liaison/consultant: Kenneth Cox, MD, MPH (USAPHC (Provisional)) Other team members: Pablo A. Aliaga, MS (Uniformed Services University of the Health Sciences); COL David M. Benedek, MD (Uniformed Services University of the Health Sciences); Susan Borja, PhD (NIMH); Tianxi Cai, ScD (Harvard School of Public Health); Laura Campbell-Sills, PhD (University of California San Diego); Carol S. Fullerton, PhD (Uniformed Services University of the Health Sciences); Nancy Gebler, MA (University of Michigan); Robert K. Gifford, PhD (Uniformed Services University of the Health Sciences); Paul E. Hurwitz, MPH (Uniformed Services University of the Health Sciences); Kevin Jensen, PhD (Yale University); Kristen Jepsen, PhD (University of California San Diego); Tzu-Cheg Kao, PhD (Uniformed Services University of the Health Sciences); Lisa Lewandowski-Romps, PhD (University of Michigan); Holly Herberman Mash, PhD (Uniformed Services University of the Health Sciences); James E. McCarroll, PhD, MPH (Uniformed Services University of the Health Sciences); Colter Mitchell, PhD (University of Michigan); James A. Naifeh, PhD (Uniformed Services University of the Health Sciences); Tsz Hin Hinz Ng, MPH (Uniformed Services University of the Health Sciences); Caroline Nievergelt, PhD (University of California San Diego); Nancy A. Sampson, BA (Harvard Medical School); CDR Patcho Santiago, MD, MPH (Uniformed Services University of the Health Sciences); Ronen Segman, MD (Hadassah University Hospital, Israel); Alan M. Zaslavsky, PhD (Harvard Medical School); and Lei Zhang, MD (Uniformed Services University of the Health Sciences).

